# Establishing a Male-Positive Genetic Sexing Strain in the Asian Malaria Vector *Anopheles stephensi*

**DOI:** 10.1101/2024.07.17.603997

**Authors:** Shih-Che Weng, Fangying Chen, Ming Li, Sammy Lee, Connor Gerry, Dylan Can Turksoy, Omar S. Akbari

**Author notes:** To whom correspondence should be addressed: Omar S. Akbari, Ph.D., School of Biological Sciences, Department of Cell and Developmental Biology, University of California, San Diego, La Jolla, CA 92093, USA. Equal contributions (Shih-Che Weng, Fangying Chen, Ming Li).

## Abstract

Genetic biocontrol interventions targeting mosquito-borne diseases require the release of male mosquitoes exclusively, as only females consume blood and transmit human pathogens. This reduces the risk of spreading pathogens while enabling effective population control. Robust sex sorting methods to enable early larval sorting in mosquitoes need to be developed to allow for scalable sex sorting for genetic biocontrol interventions. This study applies the SEPARATOR (Sexing Element Produced by Alternative RNA-splicing of A Transgenic Observable Reporter) system, previously developed for *Aedes aegypti*, to the Asian malaria vector *Anopheles stephensi*. We hypothesized that the intron from the *doublesex* gene in *Anopheles gambiae* would function in *An. stephensi* due to evolutionary conservation. Our results confirm that the splicing module from *An. gambiae* operates effectively in *An. stephensi*, demonstrating evolutionary conservation in sex-specific splicing events between these species. This system enables reliable positive male selection from first instar larval to pupal stages. RT-PCR analysis demonstrates that male-specific EGFP expression is dependent on *doublesex* sex-specific splicing events. The SEPARATOR system’s independence from sex-chromosome linkage confers resistance to meiotic recombination and chromosomal rearrangements. This approach may facilitate the mass release of males, and the cross-species portability of SEPARATOR establishes it as a valuable tool for genetic biocontrol interventions across various pest species.

## Introduction

Of all the creatures on mother Earth, the mosquito holds the grim title of the deadliest animal [1]. In the current year, a staggering 700 million cases of mosquito-borne diseases are anticipated globally, leading to over a million fatalities. *Anopheles* mosquito species stands out as they transmit malaria, one of the most widespread and lethal diseases, claiming hundreds of thousands of lives annually [2,3]. Historically, malaria control programs in Africa have predominantly targeted rural areas, where most infections occur., leading to a promising decline in the burden of malaria. However, since 2015, there have been rising concerns about malaria transmission in urban settings due to rapid urbanization, particularly in Africa [3–5]. This concern is associated with the invasion of *Anopheles stephensi*, a mosquito species notorious for its ability to transmit both *Plasmodium falciparum* and *Plasmodium vivax* malaria parasites [6–8]. Originally native to South Asia and the Arabian Peninsula, *An. stephensi* has been detected in seven African countries to date. During a malaria outbreak in Dire Dawa, Ethiopia, between April-July 2022, *Plasmodium falciparum* infections in malaria patients were strongly associated with the presence of *An. stephensi* in the household vicinity, which also tested positive for *Plasmodium* sporozoites [6–8]. Furthermore, climate change is expanding the habitable ranges of mosquitoes and their pathogens, resulting in an increasing number of people at risk of contracting mosquito-borne diseases [9–12]. Innovative technologies targeting these mosquitoes are essential for addressing this global challenge [13–16].

Genetic modification methods have been utilized in developing biocontrol strategies targeting *Anopheles* species, particularly in the context of population control [13–15,17]. Some approaches involve suppressing the female mosquito in the population since females are the primary vectors for pathogen transmission through blood meals. Methods include creating a male-biased population using gene drive technology to target essential sex determination gene *doublesex* (*dsx*) or X chromosome-linked gene [18,19]. One strategy involves the use of a drug repressible system that will kill females in the absence of the drug allowing for the release of fertile males [20]. An alternate strategy involves eliminating females through a binary CRISPR approach targeting the female-essential gene femaleless [21]. Other approaches focus on sterilizing male mosquitoes. The Sterile insect technique (SIT), for example, involves releasing sterile males released into the wild, where they mate with females but do not produce viable offspring, gradually reducing the overall mosquito population over time [22,23]. For example, the precision-guided Sterile Insect Technique (pgSIT), which enables simultaneous male selection and sterilization, ensures a release of sterilized males to suppress the population has been successfully implemented in *Anopheles gambiae* [24]. By achieving female elimination and male sterilization, these strategies aim to effectively disrupt mosquito reproduction cycles, indicating that a promising direction for biocontrol is to suppress females and select males carrying biocontrol modules, thus spreading these traits in the population. While the above methods show promise in controlling mosquito populations and reducing disease transmission, the implementation of a scalable and efficient sex-separating method is required to significantly enhance the application of these genetic strategies [25,26].

The pursuit of effective mosquito sex separation methods in *Anopheles* species has been met with progress but challenges. In *Anopheles* species, the primary method for sex separation is by observing the external morphology of the mosquitos. The differences between male and female *Anopheles* mosquitoes are subtle and challenging to discern, especially without specialized equipment or training. To address this limitation, scientists have genetically associated selectable markers with the Y chromosomes of *Anopheles* mosquitoes [27,28]. A previous study has also used testis-specific promoter-driven reporters to identify male mosquito larvae in *An. stephensi* [29]. Male-specific EGFP expression was observed as early as in approximately 45–55% of late third instar larvae, manifested as two strongly fluorescent, symmetrical ovoid bodies localized at the sixth abdominal segment. The EGFP fluorescence intensity progressively increased during larval development, allowing for the identification of male mosquitoes by the time they reached the fourth instar larval stage [29]. Alternatively, a method utilizing biased green fluorescent protein (GFP) expression to differentiate *Anopheles* larvae has been developed to *An. gambiae*, improving the efficacy of the sex-sorting marker and enabling its application as early as the first instar larval stage [30,31]. However, these methods encountered challenges as the expression patterns lacked significant divergence for effective separation, particularly during the early instar larvae stages. To better implement the genetic bio-control techniques, a reliable, scalable, and efficient sex-separating method that enables efficient early larval sex sorting is essential.

In this study, we focus on advancing the genetic biocontrol approach tailored for mosquito *An. stephensi*. Our study has developed a robust sex sorting method specifically targeting *An. stephensi*, which also holds potential for application to other *Anopheles* species. We term our method SEPARATOR (Sexing Element Produced by Alternative RNA-splicing of A Transgenic Observable Reporter). By leveraging the conserved sex-specific splicing patterns observed in *Anopheles* mosquitoes, the SEPARATOR demonstrates versatility across various species within the genus. It facilitates targeted male selection, marking a significant advancement in the field of biocontrol strategies. Furthermore, our method ensures precise and efficient sex sorting from the earliest larval stages to later developmental phases, thereby enhancing its utility for exploring biocontrol strategies tailored for *An. stephensi* and related species.

## Results

### Engineering SEPARATOR

Our previous study showcased the utilization of an engineered sex-specific spliced module derived from the *dsx* gene to produce SEPARATOR (Sexing Element Produced by Alternative RNA-splicing of A Transgenic Observable Reporter) for genetic sexing in *Aedes aegypti* [32]. *Dsx* serves as a highly conserved transcription factor pivotal in the sex determination system of insects [33–35], suggesting significant potential for application in other pest species. To enhance the applicability of SEPARATOR in genetic sexing, we conducted experiments to verify the cross-species sex-specific RNA splicing activity of the sex-specific splicing module of *An. gambiae dsx* (*AngDsx*) in Asian malaria vector, *An. stephensi*. The splicing module of *dsx* shows evolutionary conservation between *An. gambiae* and *An. stephensi*, although it is not identical (**Fig. S1**). The sex-specific splicing elements from *AngDsx* were used to construct a male-specific splicing module, designed to regulate the expression of the reporter gene (enhanced green fluorescent protein, EGFP) exclusively in males, while transgenic reporter (DsRed) expression was designed to be expressed in both sexes of transgenic mosquitoes for selection purposes. Transcriptional activation was controlled by a constitutive *Hr5IE1* AcMNPV baculovirus promoter, known for its effectiveness across various species [36–43]. The reading frame was initialized by incorporating a start codon along with a Kozak sequence, aligned in-frame with the DsRed coding sequence within the transgenic mosquito. Translation of DsRed halts due to the stop codon in the female-specific exon, regulated by female-specific RNA splicing in females. In male mosquitoes, the stop codon within the female-specific exon will be spliced out, allowing the DsRed coding sequence to align in-frame with the EGFP coding sequence. This design activates male-specific EGFP expression, facilitated by male-specific RNA splicing (**Fig. 1a**).

**Figure 1.**
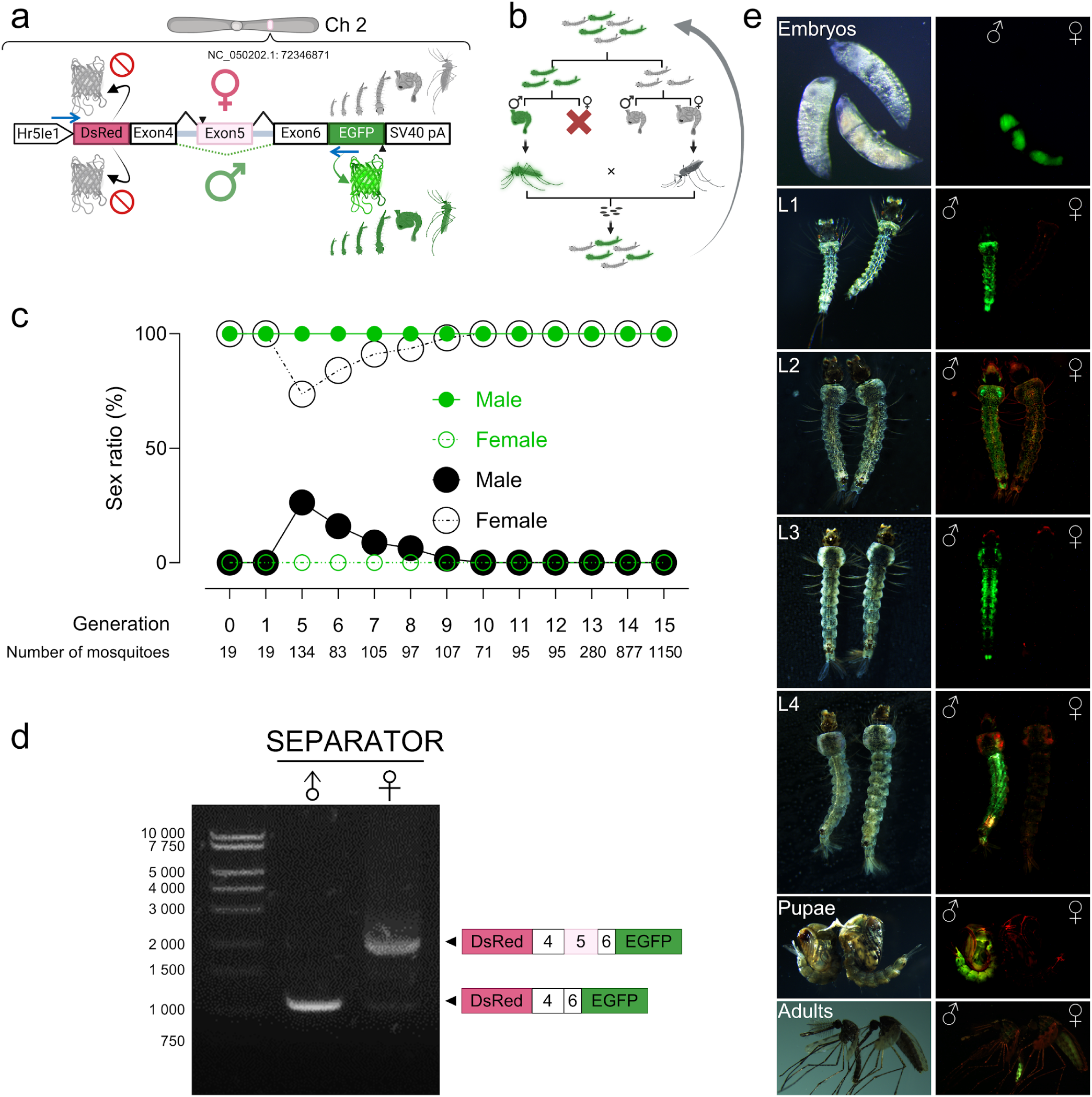
SEPARATOR in *An. Stephensi*. (a) The sex-specific splicing module of *An. gambiae dsx* (*AngDsx*) was used to construct SEPARATOR in *An. stephensi.* The expression of SEPARATOR was regulated by the *Hr5Ie1* promoter, which is a constitutive promoter derived from the baculovirus. In males, the splicing process generated a specific product that was compatible with the EGFP coding sequence, the EGFP protein. However, to ensure sex-specific expression, stop codons were strategically kept in exon 5, effectively preventing the in-frame expression of EGFP in female organisms. The SV40 pA sequence played a crucial role as the polyadenylation signal, ensuring the proper termination and processing of the SEPARATOR transcript. The blue arrows indicate the relative positions of the primer target sites used for RT-PCR. The construct is not to scale. (b) EGFP-positive mosquitoes are crossed with EGFP-negative female mosquitoes to maintain and increase the percentage of homozygous offspring. (c) Sex ratios were determined within GFP-positive and GFP-negative larvae across generations(GFP-positives are: in green symbol;GFP-negatives are in black). Sex was determined by examining the morphological differences in genital lobe shape at the pupal stage using a microscope, which are specific to each sex (male: closed circle;female: open circle). (d) Sex-specific RNA splicing of SEPARATOR. Ten EGFP-positive and ten EGFP-negative mosquitoes at the pupal stage were used. Subsequently, total RNA was extracted from each group. To investigate the splicing patterns, RT-PCR was conducted using specific primers that targeted the 3’ end of the *Hr5Ie1* promoter sequence and the 5’ end of the EGFP coding sequence. The PCR products underwent agarose gel electrophoresis, followed by gel purification and sequencing to confirm the splicing patterns. The resulting splicing patterns are shown in the right panel. (e) The developmental stages of SEPARATOR mosquitoes, including embryo, larva, pupa, and adult, were imaged using a fluorescent stereomicroscope (Leica M165FC). Eggs aged 24–48 hours post-laying were treated with a 30% NaOCl solution (approximately 3.6% active chlorine at the final concentration) for 15–30 minutes to remove the chorion and visualize the embryo. At 48-72 hours post-laying, the unhatched eggs were transferred to deionized water to collect the nascent L1 larvae. Larvae aged 0-3 hours post-hatching were collected as L1 larvae. The images are presented in two panels: the upper panel displays bright-field images, while the lower panel shows the GFP/mCH channel images.

The SEPARATOR construct was introduced into the mosquito genome to establish a genetic sex-sorting strain using the piggyBac transposon. The initial objective was to ensure that all mosquitoes expressing EGFP would be male, while DsRed would serve as a dominant marker for transgenic mosquitoes, expressed in both sexes (**Fig. 1a**). Interestingly, following microinjection, three EGFP-expressing larvae were observed, all identified as male during the pupal stage in G0, but no DsRed-expressing larvae were observed in G0. Subsequently, all pupae from G0 were sexed, and resulting adults were crossed with wild-type mosquitoes to establish stable transgenic lines based on fluorescence markers in G1. From G1 onwards, consistent results were observed, with 100% of the whole body EGFP-expressing larvae being male, and no DsRed-expressing larvae were detected.

### Precision sorting of male mosquitoes with SEPARATOR

To maintain SEPARATOR transgenic lines and enrich the population of homozygotes, larvae were sorted based on fluorescence into EGFP-positive and EGFP-negative groups. The sex in both groups was then examined at the pupal/adult stage. EGFP-positive males were subsequently crossed with EGFP-negative females (**Fig. 1b**). Over the course of 15 generations, a total of 1556 EGFP-positive larvae were manually screened, and their resulting sex at the pupal and adult stages was confirmed. Remarkably, 100% of the EGFP-positive larvae were male (**Fig. 1c, Table S1**). To verify that EGFP-expressing mosquitoes are genetic males at the larval stage, the Y chromosome-linked gene (AnSteYg) was used as an indicator [44]. Larvae from generation 9 were sorted into EGFP-positive and EGFP-negative groups based on fluorescence. Genomic DNA was extracted from each larva and subjected to PCR for Y chromosome detection. Most EGFP-negative larvae, which lacked Y chromosomes, were transgenic females. The remaining EGFP-negative larvae, which had Y chromosomes, were non-transgenic males, indicating that this generation was not completely homozygous. In contrast, all EGFP-positive larvae possessed the Y chromosome, confirming them as transgenic males. Our results showed that within the heterogeneous population, all EGFP-positive larvae were transgenic males (**Fig. 1c** and **Fig. S2**). Given that precise male mosquito selection is crucial for diverse mosquito control strategies [45–49], this showcases the robustness of our positive male selection system. To further assess the utility and robustness of SEPARATOR throughout the mosquito life cycle, we investigated the timing of EGFP expression. Our observations revealed robust EGFP signals from the late embryo stage to adulthood in mosquitoes (**Fig. 1e**). These results demonstrated that the SEPARATOR mosquitoes are sufficiently able to sort out male mosquitoes from the early stages of the mosquito life cycle.

### Male-specific gene expression regulated by sex-specific alternative RNA splicing

To validate the sex-specific splicing pattern of the SEPARATOR derived from the *AngDsx* splicing module in *An. stephensi*, a comprehensive analysis was conducted using reverse transcription polymerase chain reaction (RT-PCR) and sequencing. Ten EGFP-positive pupae and ten EGFP-negative pupae were collected for further analysis. RT-PCR was performed using primers designed to target the 3’ end of the *Hr5IE1* promoter and the 5’ end of the EGFP sequence. Subsequently, DNA sequencing was carried out to analyze the PCR products obtained (**Fig. 1d**). The sequencing result aligns with the predicted sex-specific RNA splicing patterns (**Fig. S4**). Our findings demonstrated that the sex-specific splicing module of *An. gambiae dsx* exhibited sex-specificity in *An. stephensi*. By examining the sex-specific RNA spliced transcripts, both male and female transcripts containing DsRed coding sequences were detected, in-frame with the first start codon of the splicing module, as expected. This indicates that the splicing pattern of female-specific products does not hinder DsRed expression. Nonetheless, DsRed signals were not detected in the transgenic larvae (**Fig. 1c** and **Fig. 1e**). To confirm that male-specific EGFP expression does not result from sex chromosome linkage, we determine the transgene integration site. We extracted genomic DNA from EGFP-positive mosquitoes and performed inverse PCR. Our sequencing analysis unveiled that the SEPARATOR construct had been integrated into autosome [50], specifically chromosome 2), where it intersects with the ASTEI20_038714 gene, identified as the 60S acidic ribosomal protein P0 (**Fig. S3**). Taken together, our results suggest that the expression of male-specific EGFP originating from SEPARATOR is attributed to male-specific RNA splicing.

## Discussion

To be compatible with vector control strategies aimed at combating *Anopheles* mosquito-borne diseases, we utilized the early-stage sex sorter approach SEPARATOR for selecting transgenic males. This system derives alternative RNA splicing of the sex determination gene *dsx* in *An. gambiae* and selects males by recognizing positive EGFP expression for *An. stephensi*.

We successfully implemented the sex-specific *dsx* module from *An. gambiae* in *An. stephensi*. Utilizing this module offers several advantages. Firstly, the male-specific EGFP expression is not linked to the Y chromosome, which circumvents chromosomal recombination events typically observed during large-scale insect rearing and prevents disruption of any modules linked to the sex chromosomes [26,51,52]. Secondly, the *dsx* gene is integral to the broader sex determination pathway and is expressed in both males and females across various body tissues [33,53–55]. This widespread expression allows the module to function in multiple tissues and, when combined with ubiquitous promoters, enables expression in diverse tissues throughout developmental stages. Our strategy is not limited to specific tissues, which enhances the efficiency of fluorescence sorting at early developmental stages of mosquitoes. Ultimately, the successful application of the *dsx* module across species demonstrates its potential for transferability among different species. This success can be attributed to the evolutionary conservation of the *dsx* gene sequence and function.

By sequencing the female-specific mRNA splicing fragment, our results notably demonstrate that the female-specific exon 5 in SEPARATOR is shorter than the corresponding exon in *An. gambiae* [56]. This finding suggests that the female-specific RNA splicing of SEPARATOR in *An. stephensi* favors the splicing acceptor site within exon 5 of *An. gambiae*, rather than the expected splicing acceptor site located in the intron region [56]. Bioinformatic analysis on VectorBase detected two female exon-containing splicing products by comparing RNA sequencing data against whole-genome sequencing data in *An. stephensi* [57,58]. Our RT-PCR result with wild-type *An. stephensi* corroborates the bioinformatics analysis (**Fig. S5**), demonstrating the presence of two female exon-containing transcripts resulting from alternative 3ʹ splice-site selection on the female-specific exon (exon 5). One product contains the full-length exon 5, while the other contains a truncated version of exon 5, resulting in different C-terminal ends of the female-specific DSX protein. In *Ae. aegypti*, more than one female-specific transcript is produced by the female exons (exon 5a and exon 5b) selected through exon skipping [33]. This difference suggests a divergence in the evolution of sex-determination mechanisms within Diptera. However, the regulatory mechanisms and functional implications of the two splicing products have yet to be elucidated. Further investigation is warranted to reveal the alternative splicing mechanism of the *dsx* gene and enhance our understanding of sex determination in *Anopheles* mosquitoes.

Unexpected results were observed with the SEPARATOR construct, which was designed to facilitate DsRed expression in both male and female *An. stephensi*. However, no DsRed-positive mosquitoes were detected. Despite the *dsx* splicing module in SEPARATOR generating the expected sex-specific spliced transcripts and preserving the DsRed coding sequence in both male- and female-specific transcripts, no DsRed signals were observed in either sex. This might result from improper folding or functionality of the DsRed protein due to the fusion with the DSX peptide altering its structure or interfering with its normal folding process. As a result, while our system could precisely identify male mosquitoes at the early developmental stage, it could not reliably distinguish transgenic females from non-transgenic individuals in the heterozygous SEPARATOR mosquitoes due to the non-detectable DsRed signal. Further studies will continue to utilize SEPARATOR for precise identification of both males and females in *Anopheles* mosquito populations.

Despite challenges with female selection, the SEPARATOR ensures precise and efficient male-only selection. We confirmed 100% of the EGFP-positive *An. stephensi* larvae with the SEPARATOR were male, and the earliest stage at which selection occurs is the late embryo stage. In a previous study, testis-specific promoter-driven reporters in *An. stephensi* enabled the identification of male mosquitoes in the earliest at third instar larvae, indicated by fluorescence localized at the sixth abdominal segment [29]. Our study advances *An. stephensi* sex sorting to the 1st instar stage using the SEPARATOR. Allowing separating sex in such an early developmental stage contributes to major savings in food, space, time, and labor during mass rearing. Furthermore, by utilizing the *dsx* module, the SEPARATOR overcomes the limitations of tissue-specific promoters or methods associated with EGFP expression levels, thereby improving the efficacy of the sex-sorting marker.

Notably, SEPARATOR is a positive selection method. Previous studies have demonstrated genetic sexing strains using X chromosome-linked reporters and female-specific splicing-regulated reporter expression in *Anopheles* mosquitoes [30,31,59]. For male-only releases, the positive male selection system is more robust than negative selection sexing systems. Compared to negative selection that requires the removal of 100% females to separate males from the mosquito population (such as sorting males by using female-specific *dsx* mRNA to remove females), we positively select male mosquitoes by male-specific *dsx* mRNA. This allows for the efficient identification and retention of males, streamlining the sorting process and reducing the need for complete female elimination, and resulting in a more efficient and less labor-intensive method for obtaining male-only populations. Additionally, the positive selection method used by SEPARATOR avoids issues related to spontaneous DNA mutations that can hinder the accuracy of negative selection. Consequently, SEPARATOR can significantly enhance mosquito control by enabling the selection of male-specific traits in both homozygous and heterozygous SEPARATOR mosquitoes. This is particularly useful for strategies such as releasing only sterile males, which can be applied in both conventional SIT and pgSIT. Since binary CRISPR approaches have been extensively studied in mosquitoes, such as pgSIT and Ifegenia (Inherited Female Elimination by Genetically Encoded Nucleases to Interrupt Alleles), SEPARATOR can be used to select males for either or both the Cas9 line and sgRNA lines at the larval stage. To further automate the SEPARATOR on a large scale, integrating it with the Complex Parametric Analyzer and Sorter (COPAS, Union Biometrica) presents a promising application. The COPAS instrument, known for its ability of sexing by using dominant reporter fluorescence, has been successfully used in our previous sexing systems, including *Drosophila melanogaster* and *Ae. aegypti* [32,60]. Therefore, by fully automating the sorting process, the SEPARATOR will enhance speed, facilitating the scalability of genetic technologies.

Taken all together, the SEPARATOR system represents a significant advancement in genetic biocontrol for *Anopheles* mosquitoes. It effectively selects males through positive *dsx* mRNA-based identification, ensuring precise male selection. SEPARATOR represents a significant biocontrol advancement, ensuring accurate male selection and advancing genetic biocontrol efforts against mosquito-borne diseases.

## Materials and Methods

### Molecular Cloning and Transgenesis

Plasmids were constructed using the Gibson enzymatic assembly technique. DNA fragments were amplified from existing plasmids and genomic DNA of the *An. gambiae* G3 strain using Q5 Hotstart High-Fidelity 2X Master Mix (New England Biolabs, Cat. #M0494S). All primers (Integrated DNA Technologies) are listed in **Table S2.** The resulting plasmids were introduced into chemically competent Zymo JM109 E. coli (Zymo Research, Cat # T3005), amplified, and purified using the Zymo Research Zyppy plasmid miniprep kit (Cat. #D4036). They were then sequenced using Sanger sequencing. The chosen plasmids underwent maxi-preparation with the ZymoPURE II Plasmid Maxiprep kit (Cat. #D4202) and were sequenced extensively using Oxford Nanopore Sequencing at Primordium Labs (https://www.primordiumlabs.com). The SEPARATOR plasmid (Vector 1174K), derived from the *AngDsx* splicing module, and its annotated DNA sequence map are available at Addgene (ID: 221017).

Transgenic lines were created by injecting preblastoderm stage embryos with a mixture of the piggyBac plasmid and a transposase helper plasmid. The injected G0 embryos were left to melanize for two more days before being floated in trays. Surviving pupae were sorted by sex. Male and female pupae were placed in separate cages, with a 5:1 ratio of wild-type males to females. A blood meal was provided after mating, and eggs were collected, aged, and hatched. Larvae with positive fluorescent markers were identified using a fluorescent stereomicroscope. To identify unique insertion events, transformants with fluorescent markers were bred with wild-type mosquitoes, creating distinct lines. To enhance the homozygous population, the SEPARATOR lines underwent approximately ten generations of sibling matings, selecting individuals with the most vibrant marker expression in each generation.

### Mosquito Rearing and Maintenance

*An. stephensi* mosquitoes of a strain (UCISS2018), which was previously used to generate the reference genome, was used in this study [50]. The mosquitoes were reared in incubators set at 28°C with 20-40% humidity and a 12-hour light/dark cycle, housed in Bugdorm cages measuring 24.5 x 24.5 x 24.5 cm. Adult mosquitoes had access to a 10% (m/V) aqueous sucrose solution ad libitum. Females were provided a blood meal by feeding on anesthetized mice for approximately 15 minutes, and oviposition substrates were introduced about three days after the blood meal. Eggs were allowed to melanize for an additional two days before being floated in trays. Larvae were reared in plastic containers (Sterilite, 34.6 x 21 x 12.4 cm, USA) containing approximately three liters of deionized water and were fed a mixture of ground TetraMin fish food and yeast powder. For genetic crosses, female virginity was ensured. Pupae were sexed and separated, relying on sex-specific morphological differences in the genital lobe shape (located at the end of the pupal abdominal segments, just below the paddles) before releasing them into cages. These general rearing procedures were consistently followed unless otherwise specified.

To increase the number of homozygotes in the SEPARATOR transgenic line, both high-intensity EGFP pupae and female EGFP-negative pupae were transferred to a cage and allowed to mate after eclosion. Female mosquitoes were provided a blood meal, and five adult females were individually transferred to egg tubes for colonization and egg collection. Eggs from each colony were hatched and reared. Colonies with a higher proportion of female EGFP negatives and male EGFP positives were selected for colonization in the subsequent generation.

### Fluorescent Sorting, Sexing and Imaging

Mosquitoes were examined and imaged using the Leica M165FC fluorescent stereomicroscope equipped with the Leica DMC2900 camera. We used a Leica DM4B upright microscope equipped with a VIEW4K camera for higher-resolution images. To distinguish between male and female pupae in mosquitoes, we observed the sex-specific morphological differences in the genital lobe shape located at the end of the pupal abdominal segments just below the paddles.

### Y chromosome-linked gene detection

To examine the sex of SEPARATOR at the larval stage, we isolated genomic DNA from individual mosquitoes in both EGFP-positive and EGFP-negative groups using the Blood & Cell Culture DNA Midi Kit (Qiagen, Cat. No./ID: 13343). Primers specific to Y chromosome-linked genes were employed in PCR to determine male mosquitoes among both EGFP-positive and EGFP-negative larvae [44].

### Determination of Genome Integration Sites

To identify the integration sites of the transgene, inverse PCR and Sanger DNA sequencing methods were employed according to previous studies with modifications [61]. Genomic DNA was isolated from 5 EGFP-positive and 5 EGFP-negative mosquitoes at the larval stage of SEPARATOR using the Blood & Cell Culture DNA Midi Kit by Qiagen (Catalog No. 13343), adhering to the provided instructions. Digestion of roughly 1 μg of SEPARATOR genomic DNA was carried out using HaeIII or TaqI-v2 enzymes. The digested samples were then purified and ligated overnight at 16 °C with T4 DNA ligase. This ligation mixture was subsequently utilized as a template for PCR amplification. Specific primers were synthesized to amplify the genomic regions flanking the transgene’s left and right arms. The PCR-amplified fragments were then purified through gel extraction and sequenced. The sequencing data were analyzed using BLAST to align with the *An. stephensi* genome [50], allowing for the precise determination of the transgene insertion sites by comparing the alignments from both arms. Finally, we designed a confirmed primer pair that anneals upstream and downstream of the insertion sites in the genome, enabling the amplification of the full length of the transgene element within the mosquito genome.

### Detection of sex-specific RNA splicing

Total RNA was extracted from 10 EGFP-positive and 10 EGFP-negative mosquitoes at the pupal stage utilizing the miRNeasy Tissue/Cells Advanced Mini Kit by Qiagen (Catalog No./ID: 217604), in accordance with the prescribed procedures of the manufacturer. The genomic DNA was removed with the aid of the genomic DNA eliminator column that comes with the kit. The reverse transcription was performed using both random primers and oligo dT primers to generate a comprehensive cDNA pool. To identify the sex-specific alternative splicing of the *AngDsx* splicing module, RT-PCR was utilized. Primers were specifically crafted to anneal to the terminal regions of the *Hr5IE1* promoter and the initial segment of the EGFP gene (**Table S2**). The PCR amplifications were then subjected to Sanger sequencing for detailed analysis.

## Statistical analysis

Statistical analysis was performed in Prism9 for Windows by GraphPad Software, LLC. At least three biological replicates were used to generate statistical means for comparisons.

## Data availability

Complete sequence maps and plasmids are deposited at Addgene.org (#221017). All data used to generate figures are provided in the Supplementary Materials/Tables. Generated transgenic lines are available upon request to O.S.A.

## Acknowledgments

We thank Judy Ishikawa, Ava Stevenson, Leyna Nguyen, Louis Chan, and Minzhou Ou for helping with mosquito husbandry. This work was supported by funding from NIH awards (R01AI151004, RO1AI148300, RO1AI175152) awarded to O.S.A. Figures were created using www.BioRender.com.

## Author Contributions

O.S.A. and S.C.W. conceptualized and designed experiments; S.C.W., F.C., M.L., S.L., C.G., D.T. performed molecular analyses, and genetic experiments; S.C.W. and F.C. analyzed and compiled the data. All authors contributed to the writing and approved the final manuscript.

## Ethical conduct of research

All animals were handled in accordance with the Guide for the Care and Use of Laboratory Animals as recommended by the National Institutes of Health and approved by the UCSD Institutional Animal Care and Use Committee (IACUC, Animal Use Protocol #S17187) and UCSD Biological Use Authorization (BUA #R2401).

## Disclosures

O.S.A. and S.W. have filed a patent on this technology. O.S.A is a founder of Agragene, Inc. and Synvect, Inc. with equity interest. The terms of this arrangement have been reviewed and approved by the University of California, San Diego in accordance with its conflict of interest policies. All other authors declare no competing interests.

## Additional file 1: Supplementary figures

Figure S1. Comparing the intron structure of *doublesex* in *An. gambiae* and *An. stephensi*

Figure S2. EGFP-positive mosquitoes are Y chromosome-containing mosquitoes.

Figure S3. The full length of SEPARATOR is inserted into the *An. stephensi* genome.

Figure S4. The sex-specific *dsx* transcripts align with sex-specific RNA splicing patterns in SEPARATOR mosquitoes.

Figure S5. Two female-specific *dsx* transcripts were observed in *An. stephensi*.

## Additional file 2: Supplementary tables

Table S1. The sex sorting of SEPARATOR mosquitoes

Table S2. Sequences of primers used in this study

## Supplementary Figure

**Figure S1.**
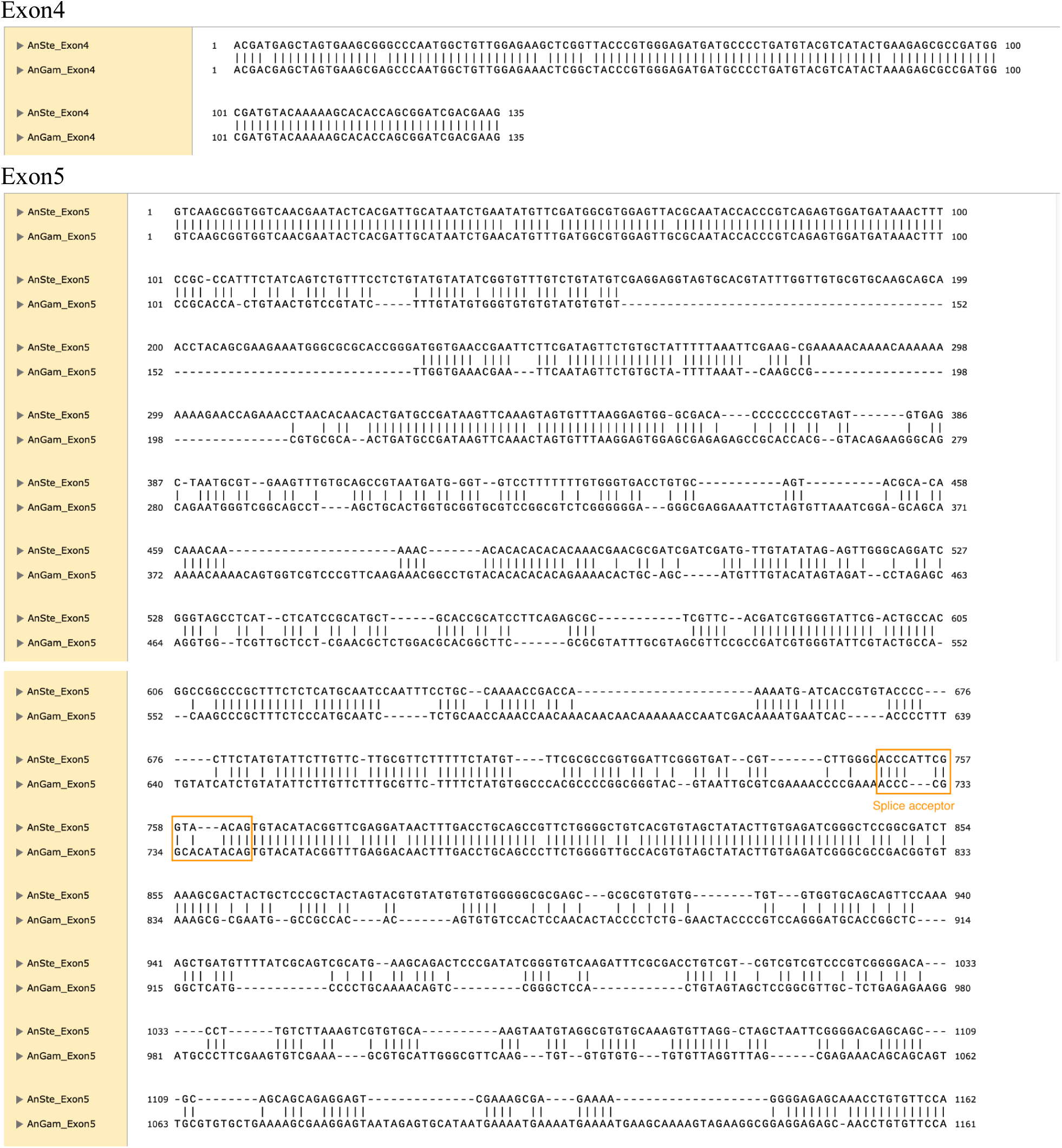
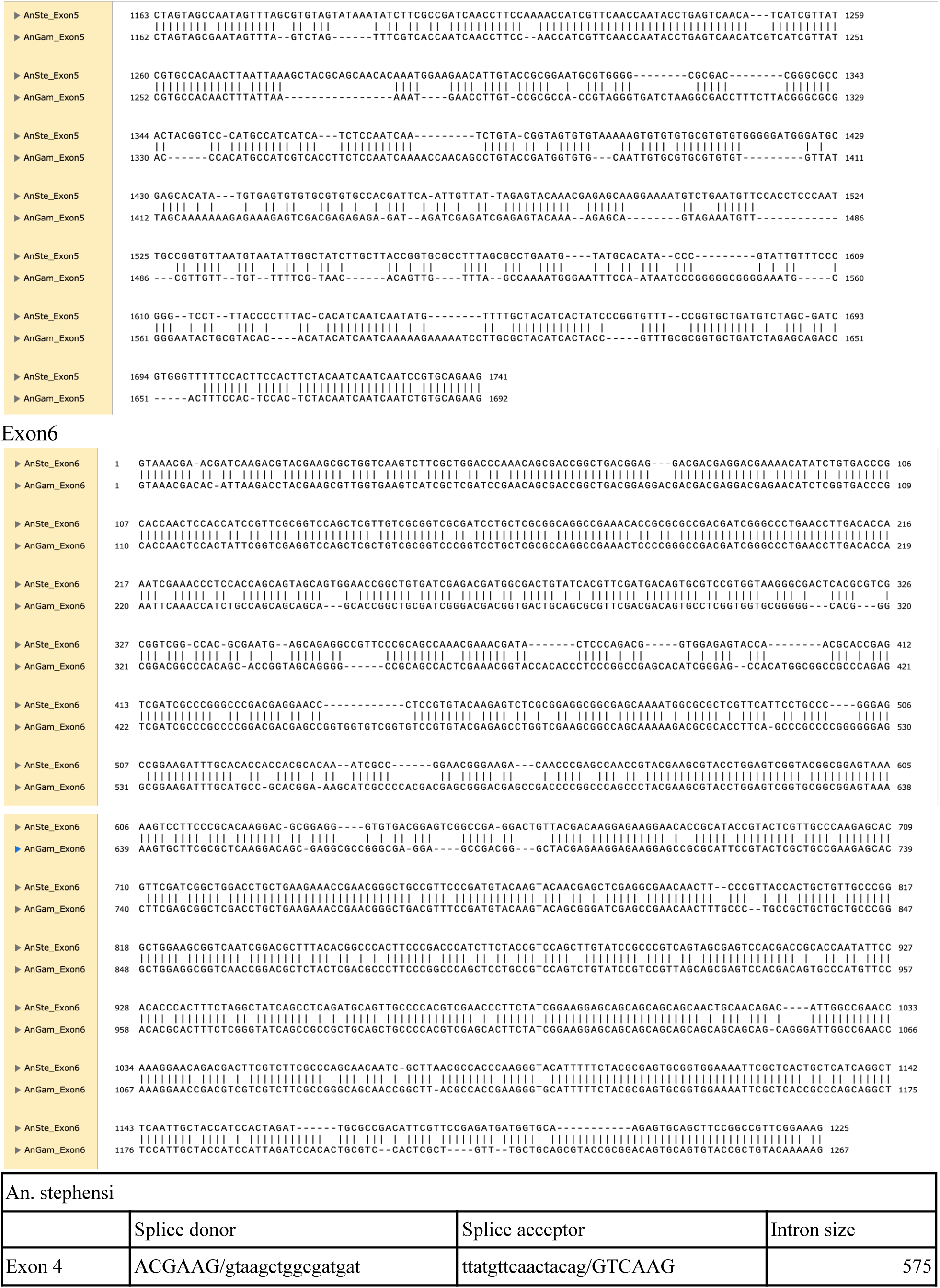
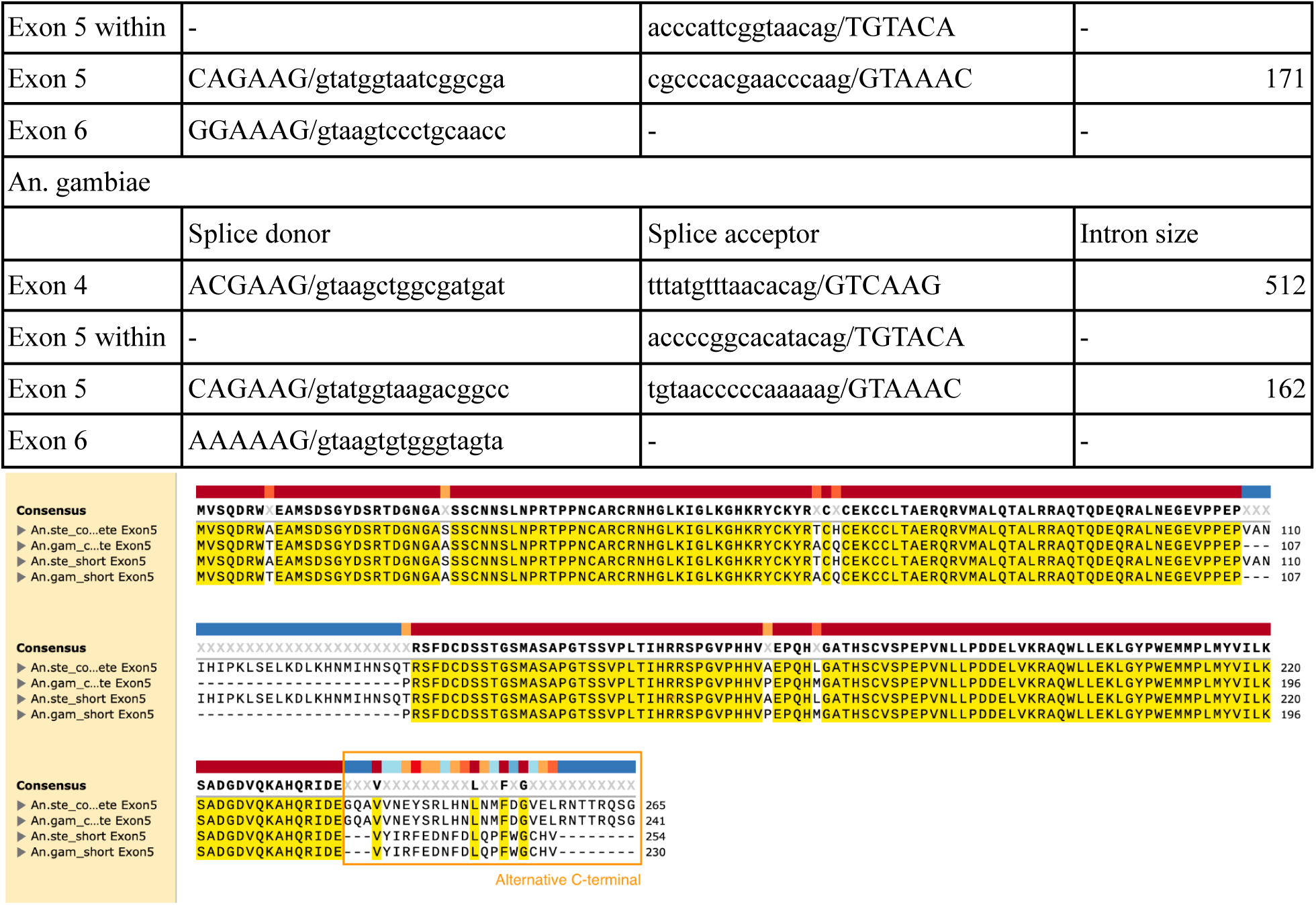
Comparing the intron structure of doublesex in *An. gambiae* and *An. Stephensi*.

**Figure S2.**
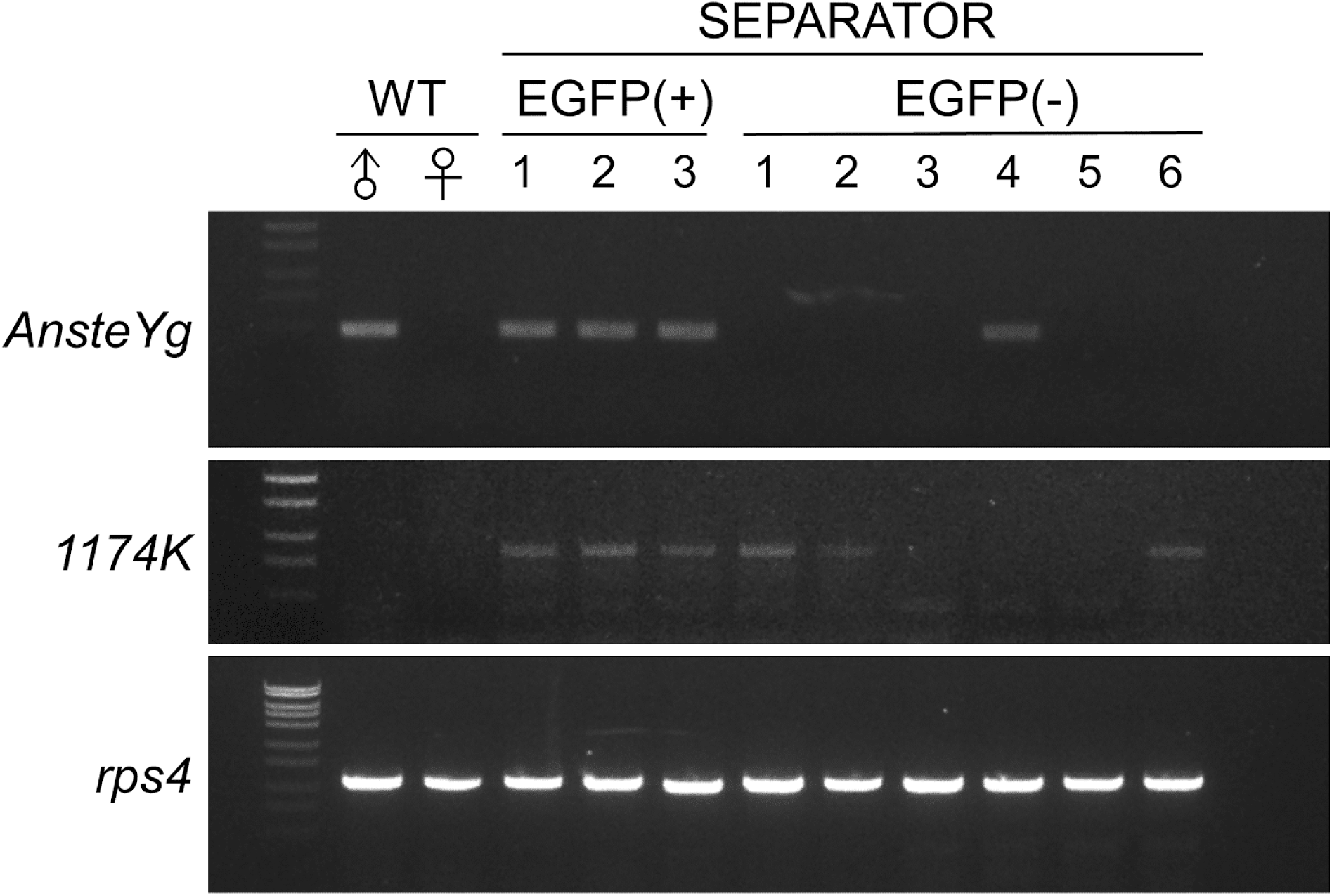
EGFP-positive mosquitoes are Y chromosome-containing mosquitoes. Genomic DNA was extracted from individual mosquitoes in both the EGFP-positive and EGFP-negative groups. PCR was performed using primers specific to Y chromosome-linked genes to identify male mosquitoes among both groups.

**Figure S3.**
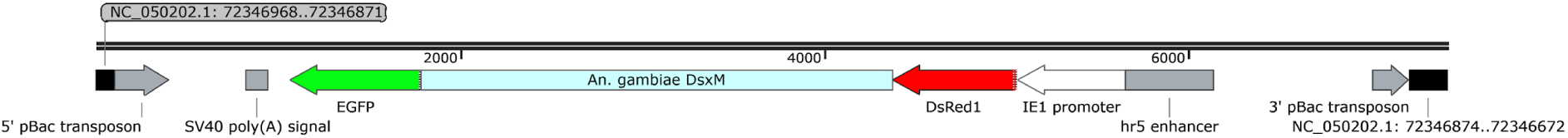
The full length of SEPARATOR is inserted into the *An. stephensi* genome. Confirmed PCR and sequencing results indicated that the full length of SEPARATOR is located on chromosome 2 (NC_050202.1: 72346871) in SEPARATOR mosquitoes.

**Figure S4.**
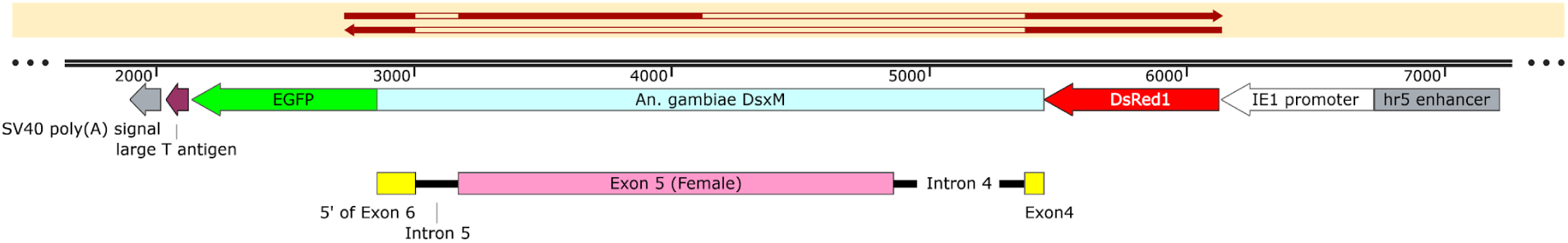
The sex-specific *dsx* transcripts align with sex-specific RNA splicing patterns in SEPARATOR mosquitoes. Both male-specific and female-specific transcripts contain the DsRed coding sequence. The female-specific exon5 has been spliced out in the male-specific *dsx* transcripts and in-frame with GFP coding sequences.

**Figure S5.**
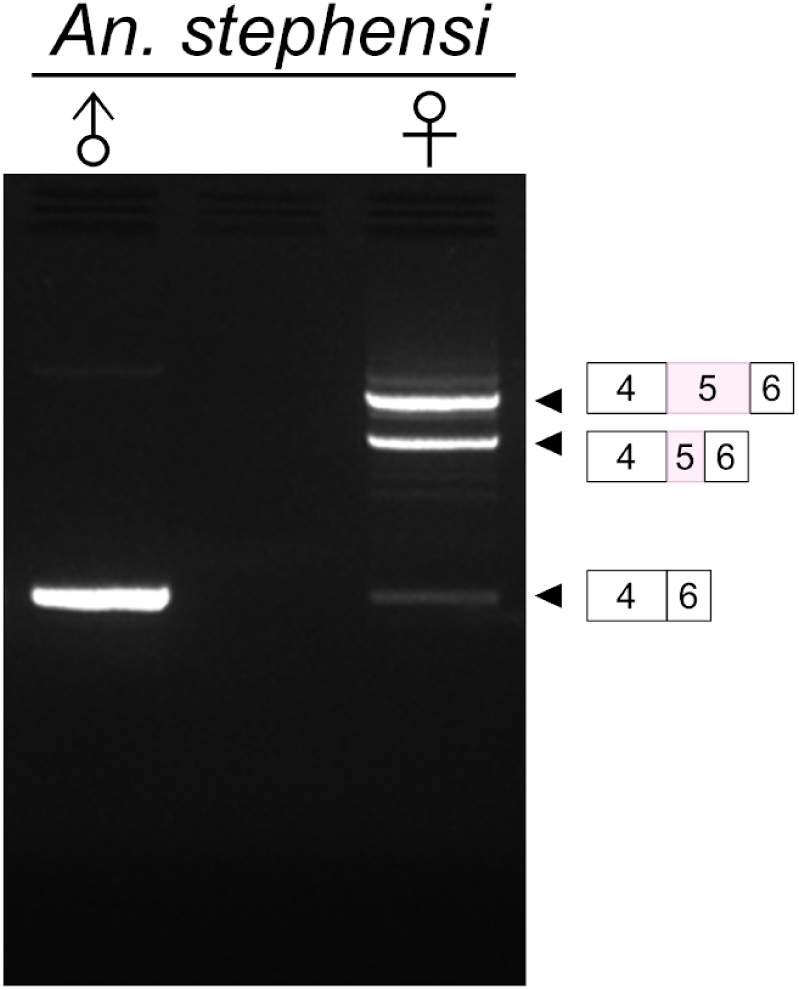
Two female-specific *dsx* transcripts were observed in *An. stephensi.* Male and female wild-type *An. stephensi* mosquitoes were selected at the pupal stage, and total RNA was extracted from each group. To investigate the splicing patterns, RT-PCR was performed using specific primers targeting the exon4 and exon6 sequences. The PCR products were then subjected to agarose gel electrophoresis, followed by gel purification and sequencing to confirm the splicing patterns. The resulting splicing patterns are shown in the right panel.

## Supplementary Tables

**Table S1.** The sex sorting of SEPARATOR mosquitoes.

**Table S2.** Sequences of primers used in this study.

